# RIL-StEp: epistasis analysis of recombinant inbred lines (RILs) reveals candidate interacting genes that control rice seed hull color

**DOI:** 10.1101/2020.06.09.141697

**Authors:** Toshiyuki Sakai, Akira Abe, Motoki Shimizu, Ryohei Terauchi

## Abstract

Studying epistatic gene interactions is important in understanding genetic architecture of complex traits in organisms. However, due to an enormous number of gene combinations to be analyzed, detection of epistatic gene-gene interactions has been computationally demanding. Here, we show a simple approach RIL-StEp, specialized to Recombinant Inbred Lines (RILs), to study epistasis using single nucleotide polymorphisms (SNPs) information of the genome. We applied the method to reveal epistasis affecting rice seed hull color phenotype, and successfully identified gene pairs that presumably control seed hull color. This method has a potential to enhancing our understanding of genetic architecture of various traits.

## Introduction

Understanding the links between genes and phenotypes of organisms is one of the most important subjects in biology. Non-additive gene interactions is called epistasis (Fisher, 1919; Phillips, 2008), and is important for crop improvement through cross breeding (Cordell, 2002; Carlborg and Haley, 2004; Xu and Crouch, 2008; Heffner *et al*., 2009; Wang *et al*., 2012).

In recent years, genome-wide association studies (GWAS) came to be widely employed to elucidate genetic variations that affect complex phenotypic traits, allowing identification of candidate loci controlling crop phenotypes (Huang *et al*., 2012; Sukumaran *et al*., 2014; Zhou *et al*., 2015). However, phenotype is affected by biological pathways that involve interactions of multiple genes (Mackay, 2014). GWAS approach has been conventionally used to identifying major Quantitative Trait Loci (QTL) associated with a phenotype of interest. In most cases, these QTL were considered as contributing additive effects to the trait values independent of the effects of other loci. If there are strong phenotypic effects of gene-gene interaction, however, GWAS approach potentially misses important loci that control the trait in combination with other loci. In such cases, additive QTL may not explain the whole phenotypic variations (Carlborg and Haley, 2004; Mackay and Moore, 2014). Therefore, it is necessary to take epistasis into account for better understanding of the genetic factors controlling phenotypic variations.

Identification of epistatic gene pairs is challenging, since one needs to consider a large number of combinations of genotypes, which incurs a heavy computational load and low statistical power due to multiple test correction. Despite these difficulties, a number of methods have been developed (Wei *et al*., 2014; Niel *et al*., 2015) that are classified into the two major approaches; the exhaustive and the non-exhaustive approach.

The exhaustive approach is designed to test all combinations of genetic variants including SNPs (Wan *et al*., 2010; Hemani *et al*., 2011; Li, 2017). The advantage of exhaustive approach is its lower risk of failure in detecting epistasis. However, the exhaustive search requires a higher computational inputs but nevertheless tends to have a lower statistical power due to multiple tests resulting from studying a large number of combinations of all pairwise genetic variations (Wei *et al*., 2014). Therefore, reduction of search space is needed to mitigate the computational burden. There have been several studies that attempted to reduce the search space by incorporating information of candidate genes based on metabolic pathways, gene ontology, and protein-protein interactions (Ritchie, 2011; Sun *et al*., 2014). However, these approaches are prone to ignoring unknown, but important, genes affecting the phenotype.

The non-exhaustive approach as represented by machine learning algorithms attempts to make non-parametric models to detect epistasis. Non-exhaustive approach is useful to detect higher-order epistatic relationships thanks to a low computational cost. However, this approach tends to generate highly complex models that sometimes suffer from a local optimality problem (Wei *et al*., 2014; Tuo, 2018). Especially when the sample size is small, the complexity of models easily becomes too large as compared to the sample size. This complexity leads to overfitting of the model to the sample dataset (Niel *et al*., 2015). Therefore, non-exhaustive approach is not appropriate in the samples with small sizes.

Recombinant Inbred Lines (RILs) are generated by first performing an intercross of genetically distinct inbred parents to obtain the F1 progeny. The F1 plants are self-pollinated to obtain the F2 plants, and each of the F2 progeny is self-pollinated several times by single seed descent (SSD) method to obtain further generations (Bailey, 1971). Each self-pollination reduces heterozygosity by half, so that after substantial number of generations (e.g. > F6), the genotypes of RILs become random mosaics of parental genotypes and the majority of genomes of RILs become homozygous. Therefore, using RILs enables one to remove the effects of heterozygous genotypes, which contributes to reducing the complexity of models used for the detection of epistasis. In addition, RILs allows phenotyping of multiple individuals from the same genotype, increasing the reliability of phenotype measurements.

In this study, we report a new approach named RIL-StEp specialized to RILs to detect epistasis in a pair of genetic variations based on the comparison of simple linear models. Bayes factor value is used to evaluating a model with epistasis against a null model without epistasis, and if this value is larger than a certain threshold, we assume that there is epistasis. This model considers the additive effects of significant QTL as well as epistatic effects between two selected SNPs. Therefore, the model is simple and easy to interpret. We applied the method to study epistatic relationships of loci that affect seed hull color of rice (*Oryza sativa*). RIL-StEp identified three candidate genes, a gene for major QTL and two epistatically interacting genes, that may control seed hull color. We suggest RIL-StEp would lead to enhancing our understanding of the genetic architecture of phenotypes of important crops as well as other organisms.

## Results

### Phenotyping of seed hull color of rice RILs

In order to quantify rice seed hull color phenotype, we converted the color to numeric values based on the CIE XYZ color space. We then measured color values of the seeds of 235 RILs of F7 generation derived from a cross between the rice cultivar “Hitomebore” (japonica type rice) and “Kaluheenati” (aus type rice). Seed hull color of RILs showed a gradation, and was not categorized into the two discrete parental phenotypes, beige and black for Hitomebore and Kaluheenati, respectively (Figure 1, Table S1). Frequency distribution of color values of the 235 RILs is skewed toward the higher phenotypic value (Figure 1); approximately one-third of RILs were whitish brown seeds (the higher phenotypic values) whereas the rest were darker brown seeds (the lower phenotypic values). From these data, we conclude that seed hull color is not controlled by a single gene, but by multiple genes.

**Figure 1.**
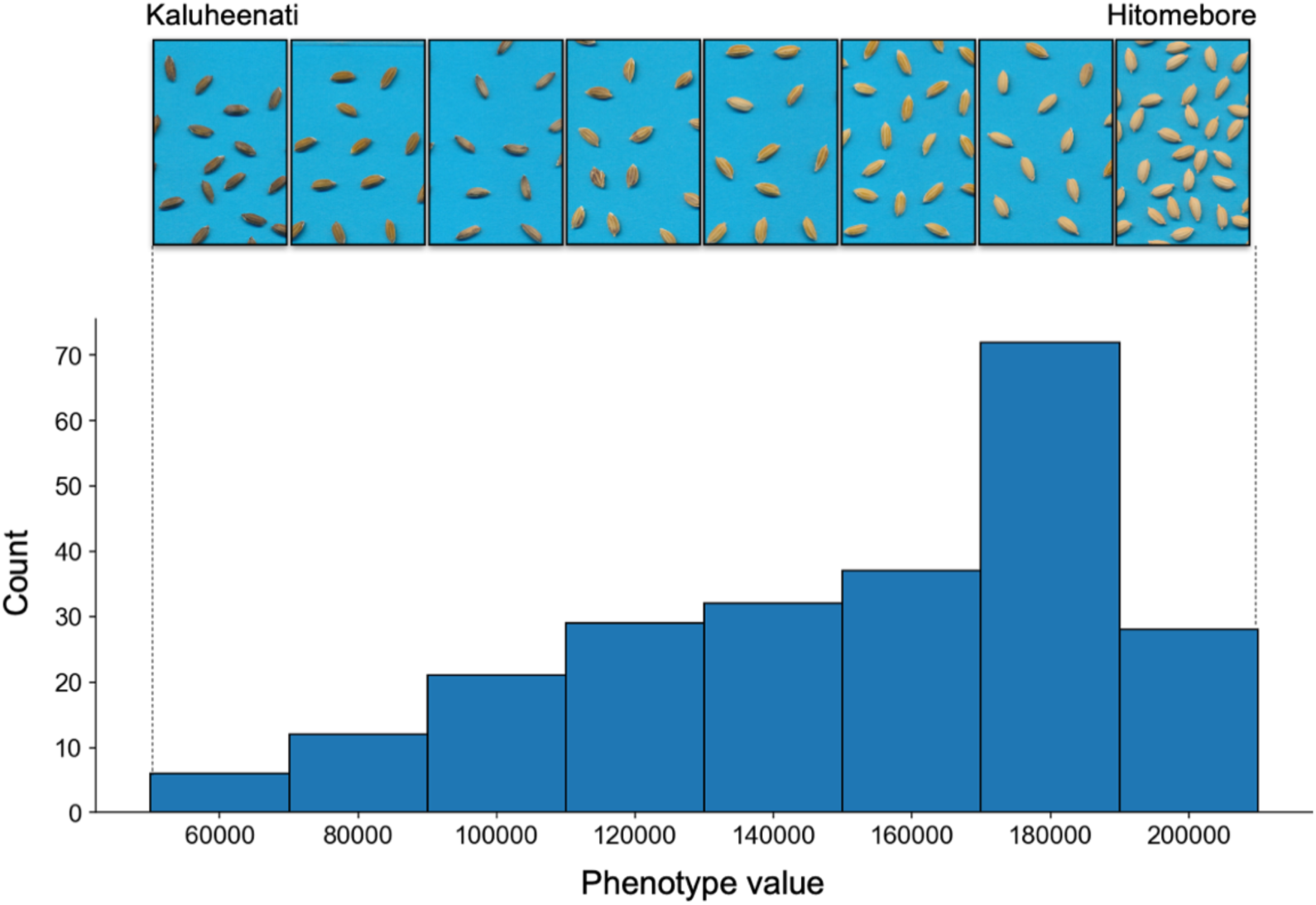
Seed hull color variation among the RILs and the distribution of phenotypic values. A histogram showing the distribution of phenotypic values. X-axis shows the range of phenotypic values of CIE XYZ color space. Y-axis shows the number of lines with phenotypic values in each range. The panels at the top show the representative images of seeds in each range of the phenotypic values.

### QTL analysis of seed hull color

We first carried out conventional QTL analysis to identify SNPs to be included in the models of RIL-StEp. Between the genomes of the two parents Hitomebore and Kaluheenati, we identified a total of 1,046,779 SNPs. We selected one SNP per 5,000bp interval and used 59,287 SNPs for subsequent QTL analysis and RIL-StEp. QTL analysis was carried out using 235 RILs by an R package “GWASpoly” (Rosyara *et al*., 2016) to detect SNPs associated with the seed hull color phenotypes. As a result, we extracted two genomic regions showing statistical significance after the Bonferroni correction, *i*.*e*. –log_10_(*p*) > 6.07 on chromosome 4 and 9 (Figure 2, Table S2). Then, we selected two SNPs showing the highest − log_10_(*p*) values in each region. These SNPs were located at chr04:23121877 and chr09: 6953870, respectively. We incorporated these two SNP values into the RIL-StEp models as the QTL variables. In order to study the possibility of epistasis of these two loci, we examined the effects of their genotypes on the phenotype. When the genotype of the SNP located in chr04:23121877 is Kaluheenati genotype, phenotype values tended to be lower (Figure S1A). The SNP located in chr09:6953870 also showed a similar tendency (Figure S1B), indicating there is no epistatic interaction between the two loci. However, when we focus on the SNP at chr04:23121877, the trait value variance of RILs with Kaluheenati genotype was larger than that of Hitomebore genotype (Figure S1C). Thus, the two QTLs do not fully explain the phenotypic variance and other genomic regions apart from these possibly affect the seed hull color.

**Figure 2.**
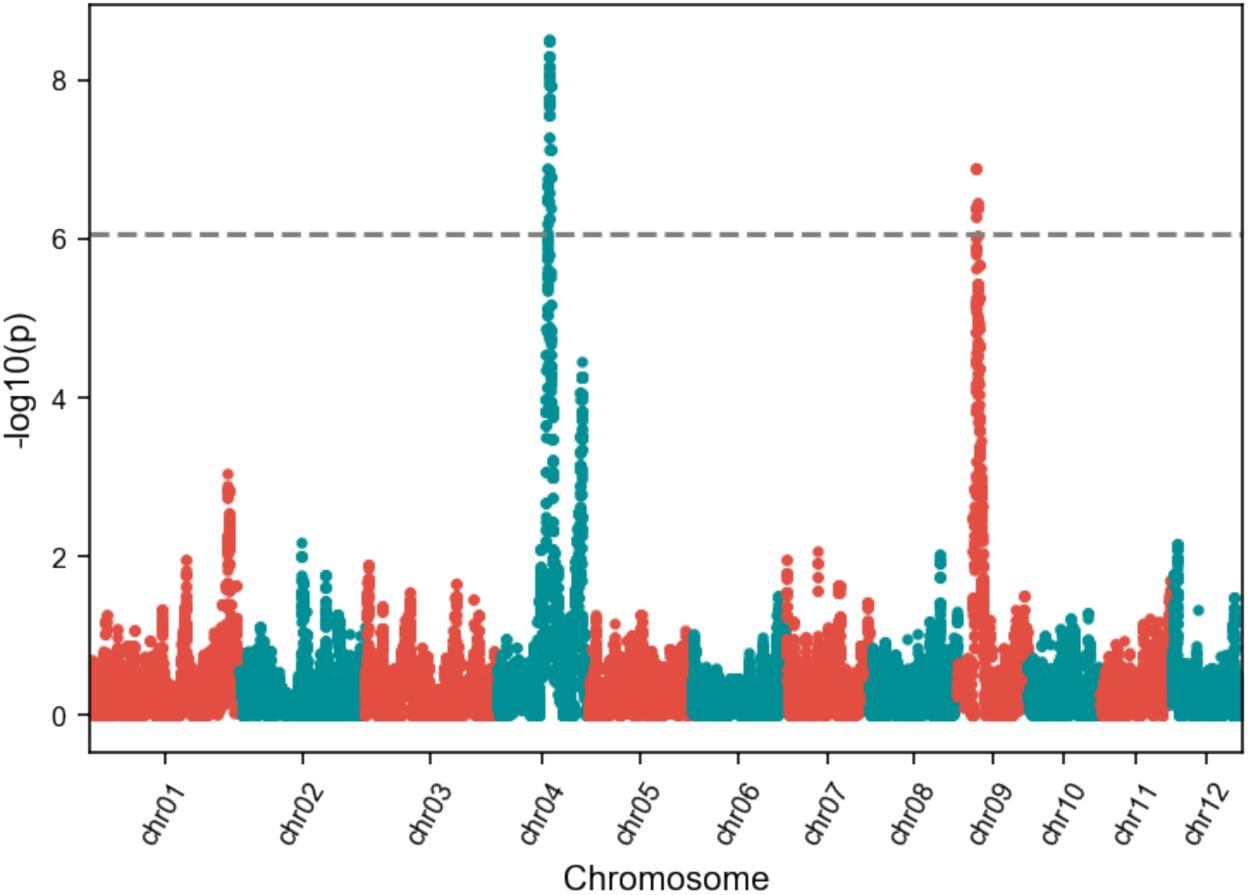
QTL analysis of rice seed hull color. A Manhattan plot showing the significant association of SNPs with seed hull color phenotype as calculated by GWASpoly (Rosyara *et al*., 2016). Y-axis shows the –log_10_(*p*) value of each SNP. X-axis shows the genomic position. Dashed line indicates the significance threshold after Bonferroni correction of multiple tests. Only SNPs located near chr04:23121877 and chr09:6953870 exceeded the threshold.

### RIL-StEp (*R*ecombinant *I*nbred *L*ines *St*epwise *Ep*istasis detection)

To detect genomic regions of RILs that are epistatically interacting, we developed a simple approach named RIL-StEp. In RIL-StEp, we generate linear models incorporating major QTLs as well as two SNPs at a time that are sampled from the entire genome. Two models, one with epistasis between the two SNPs and the other without epistasis, are compared by using Bayes factor. Specifically, we consider the following two linear models:

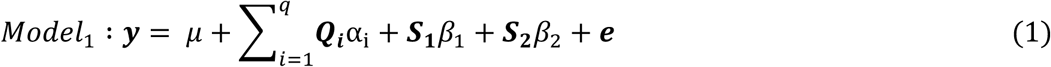

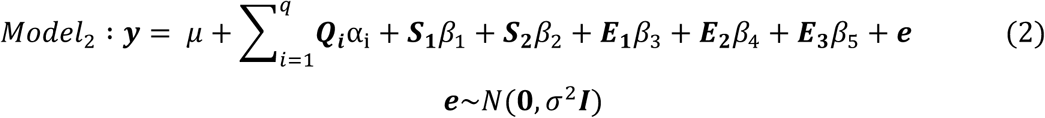

***y*** is an *n*-vector of phenotypic values for *n* samples; *μ* is an intercept term; α_i_ is the additive effect of each SNP detected by QTL analysis; *q* is the number of QTLs; *β*_1_ is the effect of first SNP and *β*_2_ is the effect of second SNP. *β*_3∼5._ are the interaction effects of the alleles from the two SNPs; *β*_3_: P1 (Parent 1) allele and P2 (Parent 2) allele; *β*_4_: P2 allele and P1 allele, *β*_5_: P2 allele and P2 allele, for the SNP1 and SNP2, respectively. One combination of alleles (P1-P1) is not included to escape multicollinearity (Table S3). ***Q***_***i***,_ ***S***_***1***,***2***_ are the *n*-dimensional genotype vectors of 1s and 0s for each QTL and the two selected SNPs. ***E***_**1∼3**_ are *n*-dimensional vectors with 1s for samples with the specific combination of alleles of selected SNPs and 0s for the rest. ***e*** is an *n*-vector of residual error and σ^2^is residual error variance.

The *Model*_1_ only includes QTLs and two selected SNPs as the variables. In the *Model*_2_, we incorporated the variables of epistasis effects between the two selected SNPs in addition to the *Model*_1_. We compared the *Model*_1_ and the *Model*_2_ based on Bayes factor. The Bayes factor is a ratio of the marginal likelihoods of the two models of hypotheses. To measure the better fit of *Model*_2_ as compared to *Model*_1_, we use the Bayes factor *K* given by:

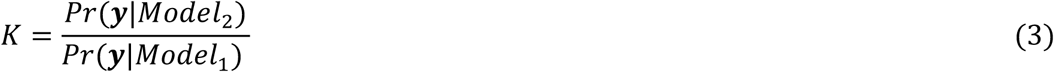

*Pr*(***y***|*Model*) is the likelihood representing the probability that phenotypic data are produced under the assumption of the *Model*. The Bayes factor *K* > 1 means the *Model*_2_ is more strongly supported by the phenotype dataset compared to *Model*_1_, indicating that the model with epistasis effects is better supported. We considered the value of *K* larger than 100 as the evidence of epistasis, following the interpretation table (Jarosz and Wiley, 2014).

### Application of RIL-StEp to rice RILs

We used RIL-StEp to detect SNP pairs showing significant genetic interactions in rice seed hull color trait. In this analysis, we incorporated two major QTLs in chromosome 4 and 9, respectively (Figure 2). To detect loci showing epistasis, we first selected one SNP every 10 SNPs out of 5,9287 SNPs across the genome, resulting in 5,929 SNPs to be considered. We applied RIL-StEp to the all pairs of the 5,929 SNPs. After calculating the Bayes factors for SNP pairs (Table S4), we focused on the genomic regions with SNP combinations showing the Bayes factor values > 100. After finding approximate positions of the loci showing possible epistasis, we applied RIL-StEp again to the combinations of all SNPs in the two regions (Figure 3, Table S5). As a result, we identified two genomic regions, chr04:22350619∼25534998 and chr04:31048756∼33482737 as the candidate regions showing epistatic interactions. The first region matched the position of the SNP detected by QTL analysis (chr04:23121877). The second region was not detected as a significant QTL; however, this corresponded to a peak with –log10p = 4.26 (Figure 2). SNP pairs between these regions showed a large Bayes factors values (Table S5). Thus, we hypothesized that the genes located in these two regions are interacting to each other. To validate this finding, we selected a SNP pair with the highest Bayes factor, and plotted the phenotype values for the combination of genotypes for the SNP pair (Figure 4). When the genotypes of SNPs located in chr04:23048862 and chr04:31581963 are both Kaluheenati types, the phenotype values tend to be low. On the other hand, if the genotypes are in other combinations, the color values were higher and similar to each other (Figure 4). This result suggested that both these regions should be Kaluheenati types to make seed hull color black. Therefore, it is assumed that two genes close to these SNPs are functioning together to determine seed hull color. To confirm this hypothesis, we surveyed candidate genes located in these two regions.

**Figure 3.**
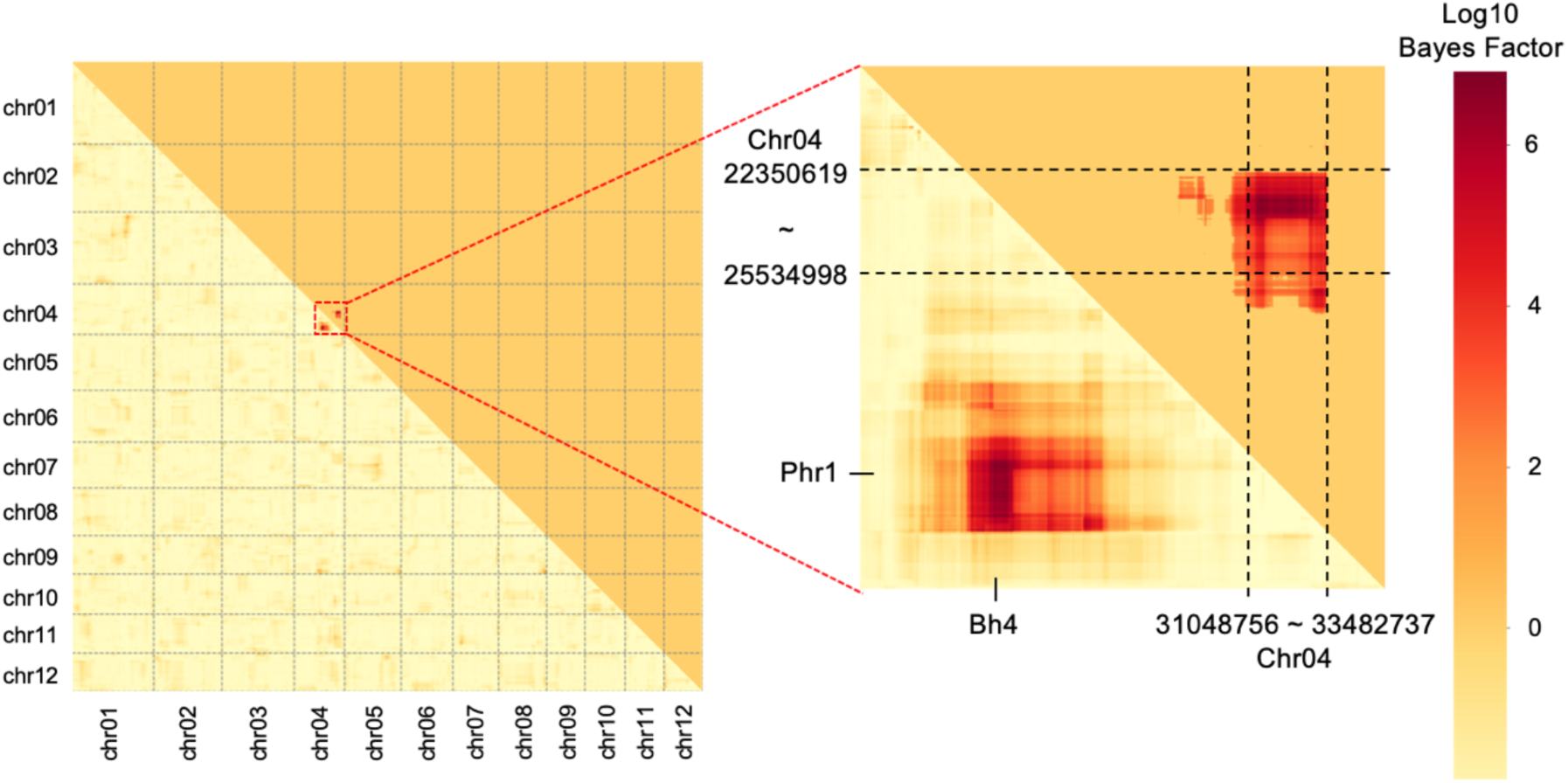
Heatmap showing Bayes factors for combinations of SNPs as revealed by RIL-StEp. The left heatmap shows the Bayes factor of SNP combinations over the whole genome. The right heatmap magnifies genomic regions with the high Bayes factors. The lower triangle shows Bayes factors of all SNP combinations. The upper triangle highlights only combinations with Bayes factors > 100. Bayes factors of all combinations of SNPs located between chr04:22350619∼25534998 and chr04:31048756∼33482737, respectively, were higher than 100.

**Figure 4.**
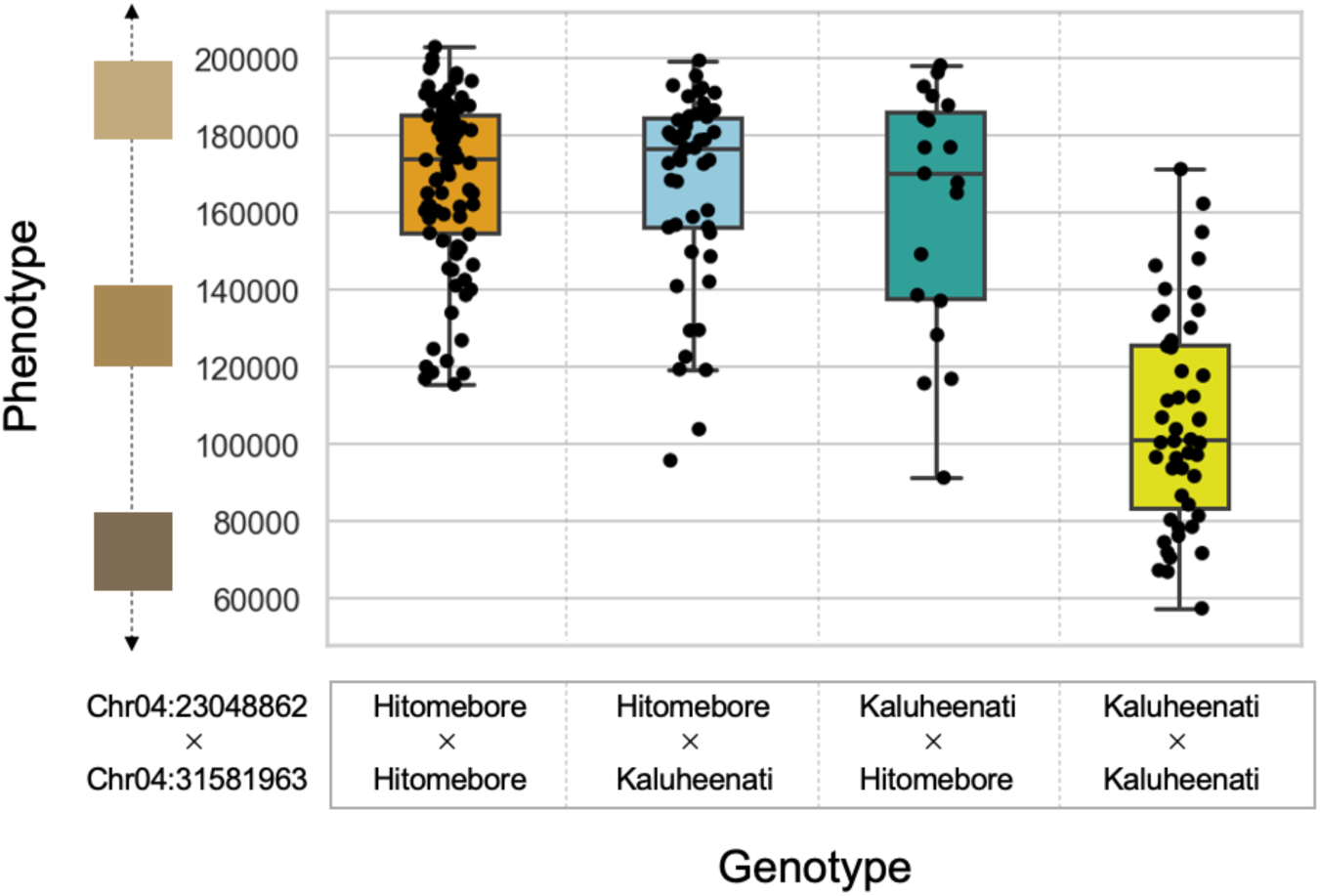
Relationships between phenotypic values and genotypes of the two epistatic SNPs as identified by RIL-StEp. A boxplot showing the phenotypic values of RILs with different combinations of genotypes of SNPs at chr04:23048862 and chr04:31581963. X-axis shows the combinations of genotypes. Y-axis shows phenotypic values. When genotypes of SNPs at chr04:23048862 and chr04:31581963 are both Kaluheenati genotype, phenotypic values tended to be low, whereas in other combinations, the values were higher and similar.

### Identifying candidate genes involved in seed hull color epistasis

We surveyed genes located in the two regions as detected by RIL-StEp and tried to identify genes that may affect seed hull color. In the region chr04:22350619∼25534998, there was *Black Hull 4* gene (*BH4* :chr04:22969845∼22971859). In the region chr04:31048756∼33482737, we found a gene called *Phenol reaction 1* (*Phr1*: chr04:31749141∼31751604). A previous study showed that the loss of function of *Bh4* changed the black hull phenotype of wild rice species to white hull of cultivated rice (Zhu *et al*., 2011). *Phr1* is known as the gene related to phenol reaction (Yu *et al*., 2008). It was reported that brown hull color of *indica* rice is caused by the presence of *Phr1* (Yu *et al*., 2008). RIL-StEp identified a pair of SNPs showing a high Bayes factor (Figure 3) and two genes close to the SNPs have been previously reported to control seed hull color. Therefore, we hypothesize that these genes are the major factors epistatically affecting seed hull color in our RILs.

We compared the nucleotide sequences of *BH4* and *Phr1* from the parental cultivars Hitomebore and Kaluheenati used for generating the RILs. Kaluheenati had intact *BH4* and *Phr1* genes, whereas Hitomebore had a 22bp deletion in *BH4* and a 18bp deletion in *Phr1* (Figure S2A, S2B). These deletions are identical to those reported in other japonica cultivars (Fukuda *et al*., 2012). In addition, these deletions were reported to cause loss-of-function in the respective genes (Yu *et al*., 2008; Zhu *et al*., 2011). Thus, we conclude that Kaluheenati maintains the function of *BH4* and *Phr1*, and Hitomebore cultivar probably lost their functions. Using a crossed line between an indica type cultivar “Habataki” and japonica type cultivar “Arroz da Terra”, Fukuda et al. (2012) reported that both *BH4* and *Phr1* are necessary for maintaining black hull phenotype. *BH4* encodes a tyrosine transporter and *Phr1* encodes a polyphenol oxidase of the tyrosinase family (Yu *et al*., 2008; Zhu *et al*., 2011). Tyrosine is converted by the tyrosinase to melanin, the main black pigment (Riley, 1997). Thus, it is assumed that *BH4* is required for transportation of tyrosine and *Phr1* for melanin biosynthesis (Figure 5). This suggests that the melanin biosynthesis pathway does not operate if either of these two genes does not function. It is consistent with the result that seed hull color tends to be beige when even one of the two SNPs is Hitomebore genotype (Figure 4).

**Figure 5.**
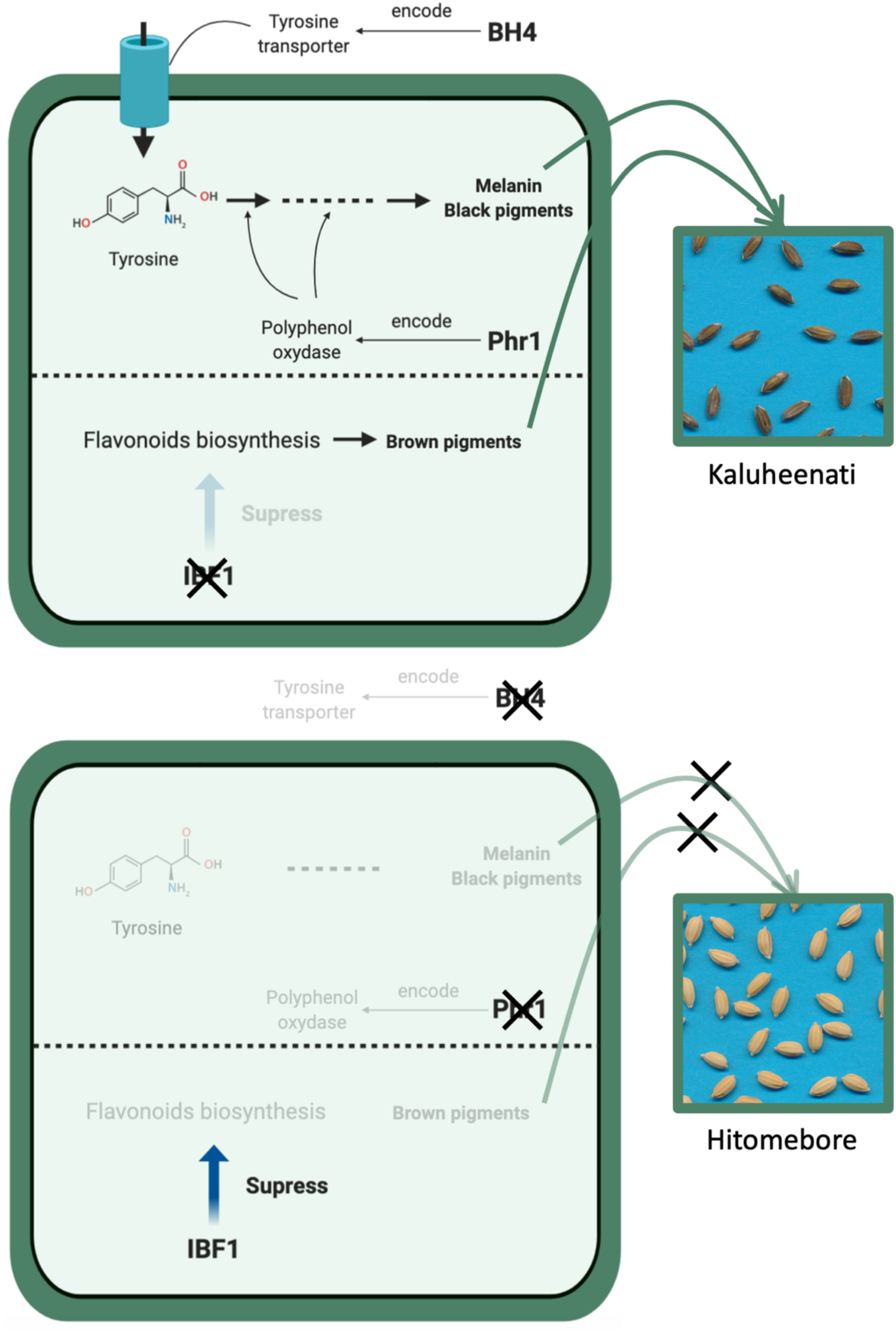
A simplified scheme of the pathways related to rice seed hull color as hypothesized in the present study. This figure shows the summary of biological function of *BH4, Phr1*, and *IBF1. BH4* encodes a tyrosine transporter (Zhu *et al*., 2011) and *Phr1* encodes a polyphenol oxidase (Yu *et al*., 2008). These genes are related to melanin biosynthesis pathway. *IBF1* inhibits flavonoids biosynthesis as a suppressor (Shao *et al*., 2012).

In addition, we surveyed genes located near the SNP chr09:6953870 as identified by the QTL analysis to address its contribution to seed hull color in combination with *BH4* and *Phr1*. We found *Inhibitor for brown furrows1* (*IBF1*) located in chr09:6873236∼6874612. The previous study showed *ibf1* mutants of japonica and indica type cultivars accumulate brown pigments during seed maturation. Thus, *IBF1* is a suppressor of brown pigment deposition in rice hull furrows (Shao *et al*., 2012). We compared the sequences of *IBF1* for the two parental cultivars. Kaluheenati had 19bp deletion in *IBF1*, whereas Hitomebore had an intact protein coding region (Figure S2C). This result suggests that 19-bp deletion in Kaluheenati caused loss of function of *IBF1*, which no more suppresses the accumulation of brown pigmentation of rice hull furrows. This is in line with the lower phenotypic value (brown color) of RILs with Kaluheenati-type genotype near the *IBF1* gene (Figure S1B). *IBF1* is reported to be involved in flavonoids biosynthesis (Shao *et al*., 2012).

Taken together, the relationship between seed hull color phenotype and genotypes of the three SNPs located near *BH4, Phr1*, and *IBF1* showed that the effect of *IBF1* is clearly independent of that of *BH4* and *Phr1* (Figure S3). Thus, the pathway involving *BH4* and *Phr1* and that of *IBF1* are probably functioning independently (Figure 5).

## Discussion

In this study, we describe a new approach “RIL-StEp” for detecting epistatic relationships of genes. This approach is specialized to RIL population and based on Bayes factors for comparison of simple linear models. Using RIL-StEp we successfully detected a likely gene pair showing epistasis that affect seed hull color. The advantage of RIL-StEp is its high interpretability as compared to other approaches that consider many variables at once. Our model includes in variables only significant QTLs as well as epistasis of two SNPs at a time without considering heterozygous genotype. Thus, our model has a low complexity without any possibility of overfitting as seen in the complex model. Therefore, we believe that RIL-StEp is a recommended option to detect epistasis in any traits when the RILs are used. Additionally, our approach adopted Bayes factors. It is known that Bayes Factors have flexibility to combine prior information on each effect of genetic variants (Wakefield, 2009; Runcie and Crawford, 2019). Thus, although we specified prior distribution according to the paper (Liang *et al*., 2008), our approach is capable to incorporate any prior information. The disadvantage of our approach is the difficulty in detecting higher-order (e.g. 3 loci) epistatic relationships. Detection of high-order relationships using our exhaustive approach increases computational cost explosively and decreases the interpretability of the models (Taylor and Ehrenreich, 2015). Therefore, the non-exhaustive approach may be more appropriate to identify the high-order epistasis.

We succeeded in identifying genomic regions that show epistasis. However, these regions contained multiple genes and we could not specify the responsible genes only by the genetic analysis. It is a challenging problems of GWAS to fill the gap between identification of the genomic regions and identification of the causative genes responsible for the phenotype (Gallagher and Chen-Plotkin, 2018). It could be possible to pin down to much smaller genomic regions by applying a more strict threshold in the epistasis analysis. However, this has a risk of missing true positive SNPs. It is known to be difficult to decide the proper balance between type I and type II errors (Todorov and Rao, 1997). Thus, the approach to identify genes using only statistical significance threshold is usually not possible and not appropriate.

In our case, we have successfully specified strong candidate genes presumably controlling the phenotype using the knowledge about the candidate genes and the sequence analysis. However, this approach may not be applicable in every case. In epistasis analysis, there may be several approaches to validate the epistatic relationship between the genes. For example, co-expression analysis explore genes in the same biological processes (Aoki *et al*., 2007; Mao and Chen, 2012; van Dam *et al*., 2018). eQTL analysis identify genetic variants regulated by specific genes (Gilad *et al*., 2008; Feltus, 2014). Therefore, combining information from other sources of evidence to RIL-StEp may enhance our capability of identifying interacting genes.

We quantitatively measured seed hull color to use as phenotypes. Seed hull color can be treated not only as categorical traits, but also as quantitative traits (Shao *et al*., 2011). In our study, the difference in seed hull color between the two parental lines is most likely controlled by the three genes. Two genes of them seem interacting to each other. Thus, seed hull color exhibited gradual change according to the genotypes of these genes.

The main factor of seed hull color is probably black and brown pigments. Thus, it is assumed that measuring seed hull colors based on the brightness in CIE XYZ color space was appropriate. However, when various colors are included, it is difficult to convert colors to 1-dimensional values and we have to consider other methods of measurement. Some of other traits are also difficult to assess, like virulence and resistance response (Stewart and McDonald, 2014; Stewart *et al*., 2016). Thus, developing methods to quantitatively measure the trait is one of the critical steps in understanding genetic architecture that affects traits.

To summarize, we propose a novel approach based on simple linear models to detect epistatic interaction for quantitative traits in the RIL population. By applying RIL-StEp to rice seed hull color, we succeeded in identifying three genomic regions related to seed hull color. Incorporating additional information allowed us to identify candidate genes involved in seed hull color variation. Thus, our approach has the potential to identify epistasis in various biological traits.

## Experimental procedures

### Materials

A rice (*Oryza sativa*) cultivar Hitomebore belonging to the japonica rice group shows white seed hull color. On the other hand, a cultivar Kaluheenati, which is one of the NARO World Rice Core Collection (WRC) (Kojima *et al*., 2005), belonging to the aus rice group shows black seed hull color (Figure 1). Hitomebore and Kaluheenati were crossed and RILs of the F9 generations consisting of 235 lines were generated by the single seed descent (SSD) method. Images of seeds of each line were scanned and saved as pictures for phenotyping of seed hull color.

## Methods

### Genotyping of RILs by whole genome resequencing

To obtain the genotypes of all RILs, we performed the whole genome resequencing of the parents and 235 RILs. We filtered and trimmed these sequences using prinseq (Schmieder and Edwards, 2011) and FaQCs (Lo and Chain, 2014). Then, the quality-trimmed Illumina short read data were aligned against the reference genome using BWA (Li and Durbin, 2009). We used genome sequence of Os-Nipponbare-Reference-IRGSP-1.0 as the reference (Kawahara *et al*., 2013). After mapping, we sorted and added index to bam files using samtools(Li *et al*., 2009). These bam files were subjected to variant calling with bcftools (Narasimhan *et al*., 2016). Finally, we imputed the variants based on Hitomebore and Kaluheenati genotypes using LB-impute (Fragoso *et al*., 2016). For biallelic SNPs in our RILs, there are three genotypic classes Hitomebore-Hitomebore, Hitomebore-Kaluheenati and Kaluheenati-Kaluheenati. These genotype classes were parameterized to {0, 1, 2}. We used 59,287 SNPs for the analysis that were selected from a total of 1,046,779 SNPs found between the two rice parents. For selection of SNP, we used only one SNP per 5,000bp interval.

### Phenotyping and quantification of seed hull color

In the RILs, seed hull color showed gradation between beige and black colors (Figure 1). It is known that quantitation of phenotypes tend to improve statistical power and interpretability of relationships between genetic variants and phenotypes (Bush and Moore, 2012). Therefore, in order to convert seed hull color phenotypes to quantitative values, we measured the brightness of the seed hull color. First, we extracted seed image from the original picture and constructed the matrix of RGB values of the image. Then, we applied Principal Component Analysis(PCA) to extract the RGB values to pick up representative color of all seeds in the image (Figure S4). We applied this process to each RIL and obtained the representative RGB value of seed hull color for each line. Finally, we converted these representative RGB values to CIE XYZ color space. Y axis value showed the brightness in CIE XYZ color space. Thus, we used y-axis values as quantitative phenotypes. The larger y-axis values express brighter color as compared to the lower y-axis values corresponding to darker color (Figure S4).

### QTL analysis

To identify SNPs corresponding to major QTLs and include these SNPs in our linear models, we used genome-wide association study (GWAS) approach based on the mixed linear model (Yu *et al*., 2006). We utilized the R package “GWASpoly” (Rosyara *et al*., 2016) to identify genomic regions that show a significant association with the phenotypic effect. We used the Bonferroni method to determine the QTL significance threshold. Then, we selected a SNP with the largest values of −*log*10*p* for each genomic region that exceeds the Bonferroni threshold. These selected representative SNPs will be included in the *Model*_1_ and *Model*_2_ as the major QTLs.

### Calculating Bayes factors in RIL-StEp

Bayes factors are computed by integrating the likelihood with respect to the priors on parameters. We estimated Bayes factors based on Monte Carlo sampling for the integration of parameters. The equation (1) and (2) can be expressed as:

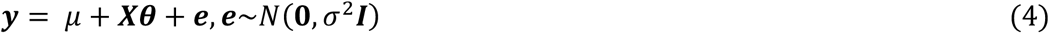

***X*** is a *n* × *r* design matrix of genotypes for QTL or epistasis variables. ***θ*** is a *r* × 1 vector of QTL and epistasis effects. *r* is sum of the number of QTLs and epistasis variables used in the model. In Monte Carlo sampling, we specified the prior distribution of ***θ*** as given by:

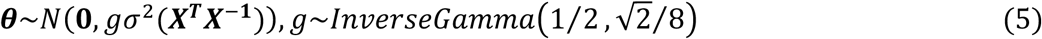

The number of iterations to estimate Bayes factor was 10,000. We used the R package “BayesFactor” (Morey *et al*., 2018) to compute Bayes factors. We applied these processes to a total of 17,573,556 combinations of SNPs. When we calculated Bayes factor, we did not consider RILs showing heterozygous genotypes at the QTL or the selected SNPs.

The source codes and detailed usage instructions of RIL-StEp are freely available from GitHub (https://github.com/slt666666/RILStEp) under MIT license.

## Data availability

The genotype dataset, seed images of RILs, and detail of supporting information (Table S4, S5) were deposited in the Zenodo (10.5281/zenodo.3882105). All other relevant data are within the paper and the supplemental files. RIL-StEp package source codes and a user manual are freely available through GitHub (https://github.com/slt666666/RILStEp) under MIT License. The scripts used in phenotyping process are also deposited in GitHub (https://github.com/slt666666/Seed_phenotyping)

## Author Contributions

R.T. and T.S. conceptualized the study. T.S., A.A., and M.S. performed research. The original draft was written by T.S., and reviewed by R.T., A.A., and M.S.

### Acknowledgements

We thank the National Agriculture and Food Research Organization (NARO) gene bank, Japan for providing the WRC seed. This study was supported by grants from the Project of the NARO Bio-oriented Technology Research Advancement Institution (Research program on development of innovative technology), and by grant JSPS KAKENHI 15H05779 and 20H00421 to RT, 17H03752 and 20H02962 to AA. We thank Sophien Kamoun for valuable comments. The authors declare that there are no conflicts of interest.

## Short legends for Supporting Information

**Table S1 Phenotypic values of seed hull color**.

**Table S2 The results of GWAS analysis using GWASpoly**.

**Table S3 Values of RIL-StEp coefficients used for each genotype**.

**Table S4 Results of RIL-StEp for combinations in whole genome regions**.

**Table S5 Results of RIL-StEp for combinations in strong candidate genomic regions**.

## Supporting Information

**Figure S1.**
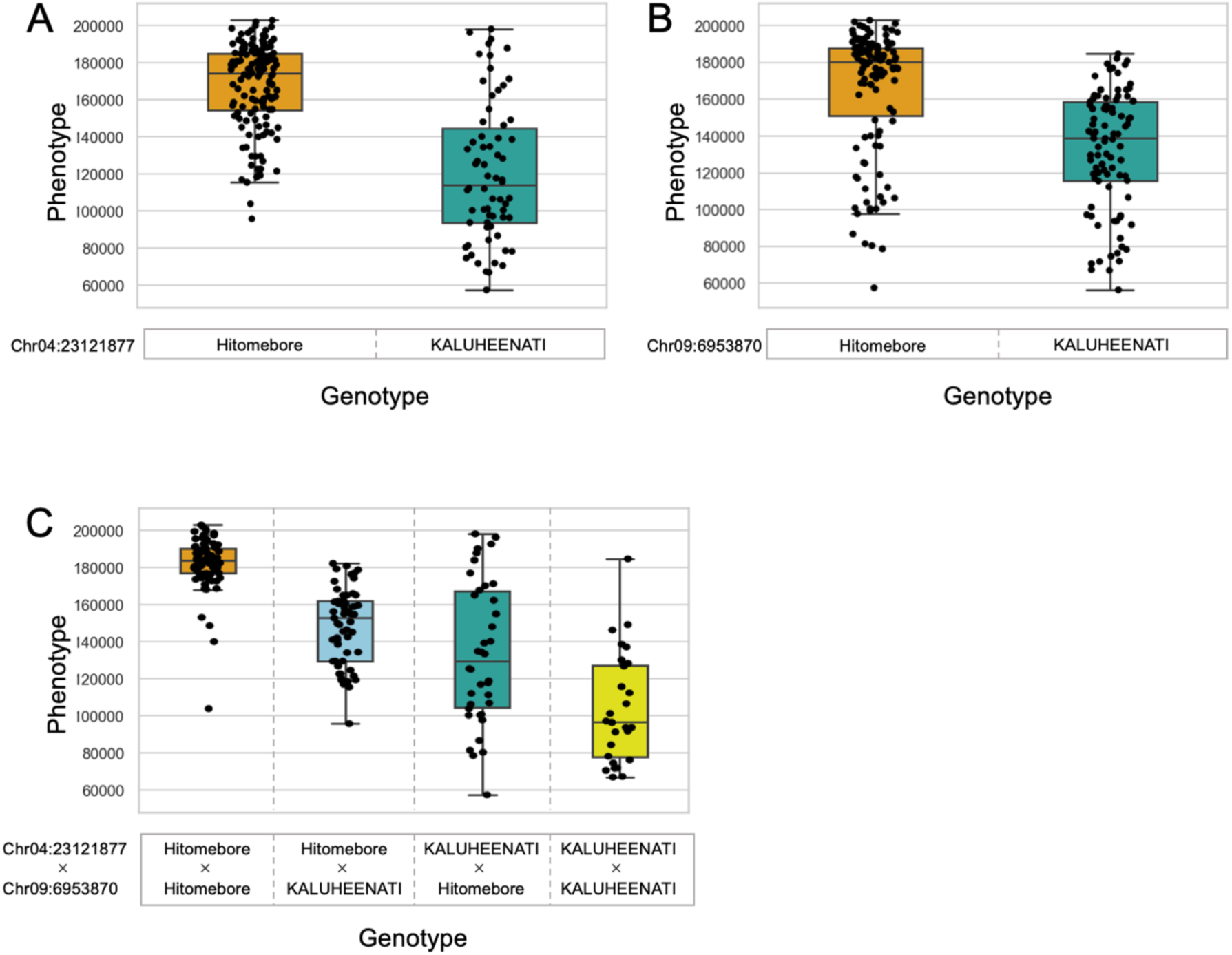
Relationships between rice seed hull color phenotypes and the genotypes of the two major QTLs. Boxplots showing the phenotypic values of RILs (Y-axis) in relation to their genotypes at the two major QTLs (Y-axis). (A) The phenotypic values of RILs separately shown for Hitomebore or Kaluheenati genotypes at the SNP of chr04:23121877. (B) The phenotypic values of RILs separately shown for Hitomebore or Kaluheenati genotypes at the SNP of chr09:6953870. (C) The phenotypic values of RILs shown separately for the four combinations of genotypes at the SNPs of chr04:23121877 and chr09:6953870.

**Figure S2A.**
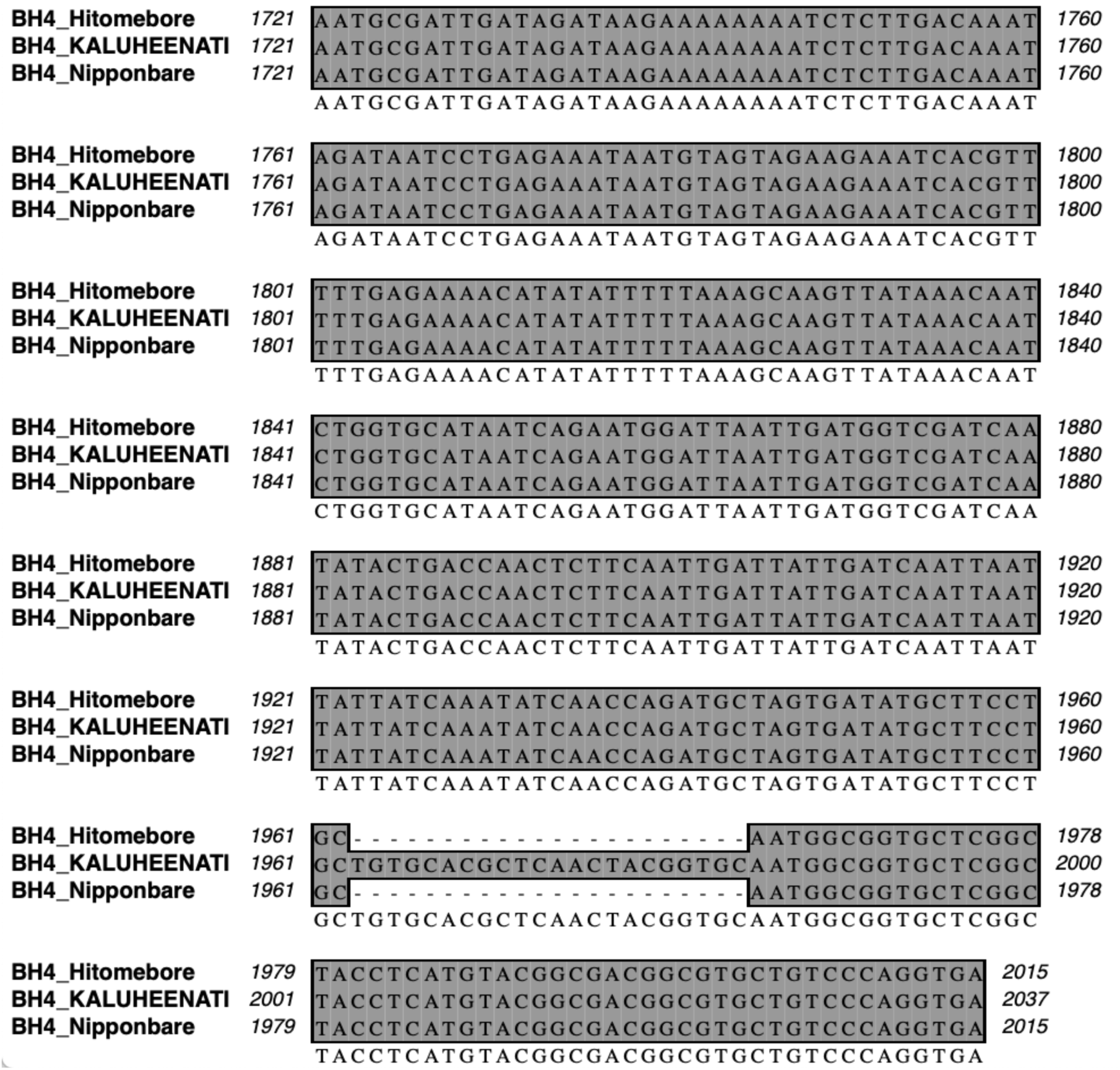
DNA sequence alignment of the *BH4* gene. Genome sequences of the *BH4* gene for Hitomebore, KALUHEENATI, and Nipponbare are aligned and shown. Hitomebore and Nipponbare sequences have a 22bp deletion.

**Figure S2B.**
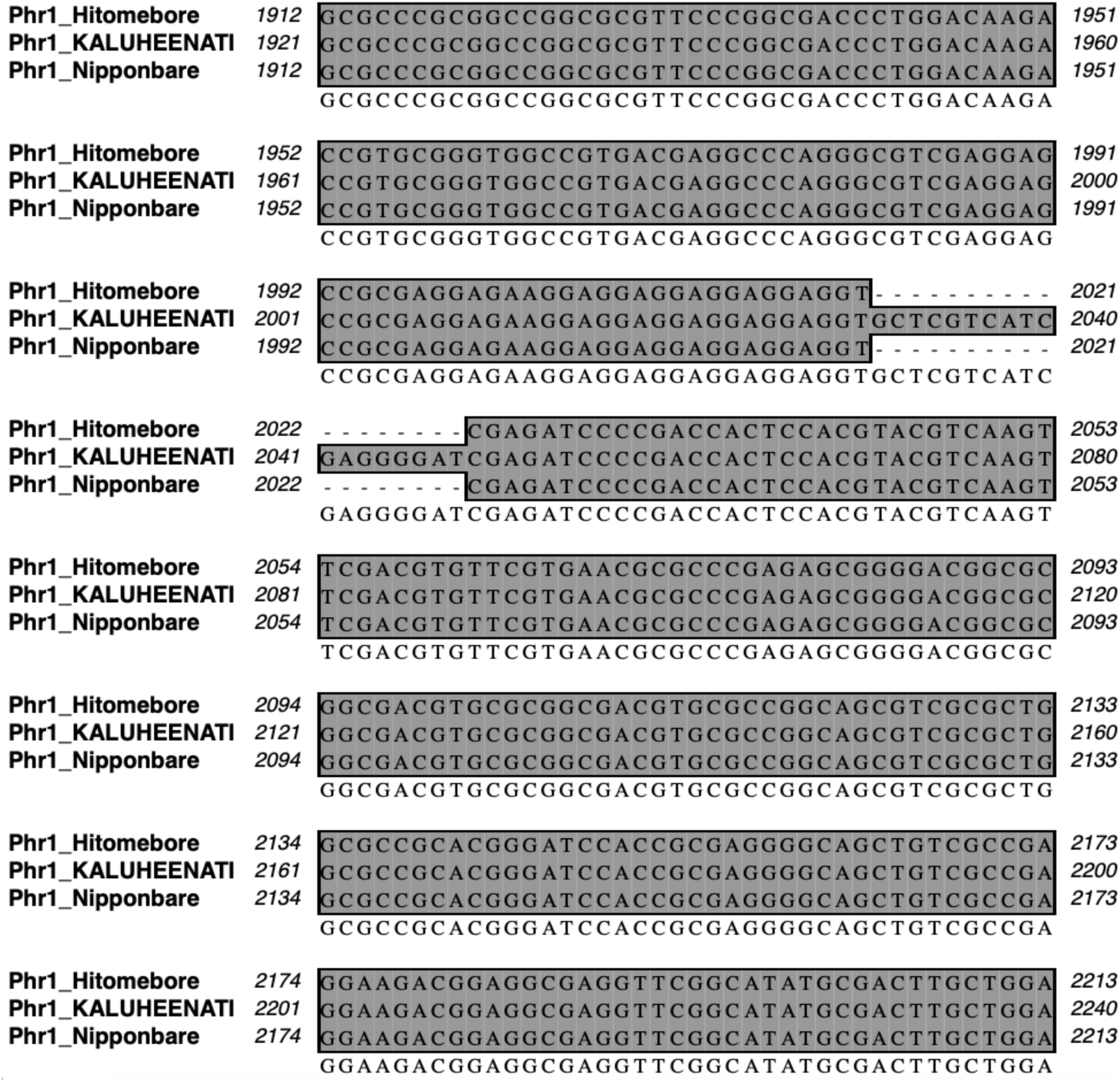
The sequence alignments of *Phr1*. Genome sequences of Hitomebore, KALUHEENATI, and Nipponbare in the region of *Phr1* gene. Hitomebore and Nipponbare have 18bp deletion.

**Figure S2C.**
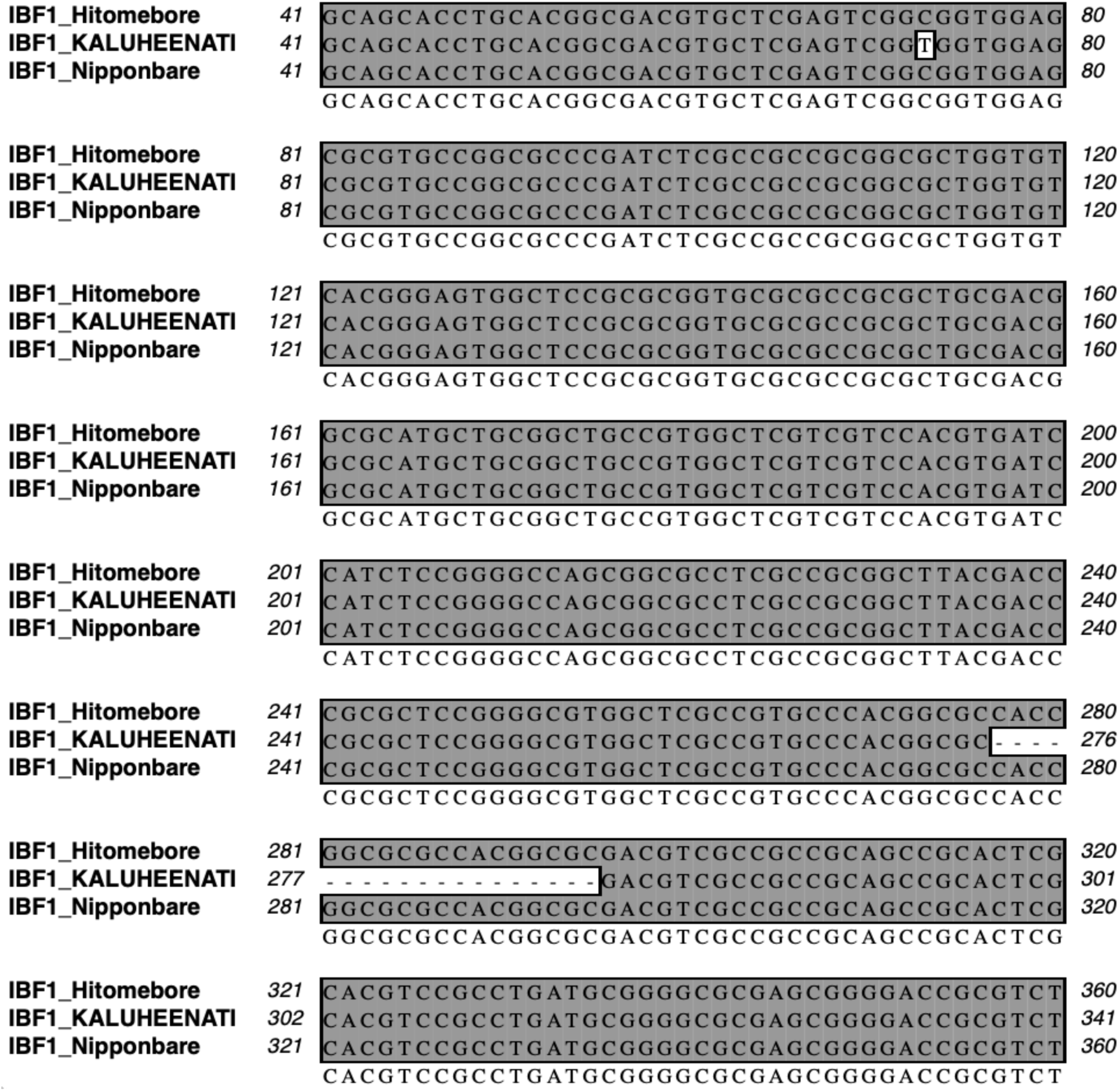
The sequence alignments of *IBF1*. Genome sequences of Hitomebore, KALUHEENATI, and Nipponbare in the region of *IBF1* gene. KALUHEENATI has 19bp deletion and 1 genetic variant.

**Figure S3.**
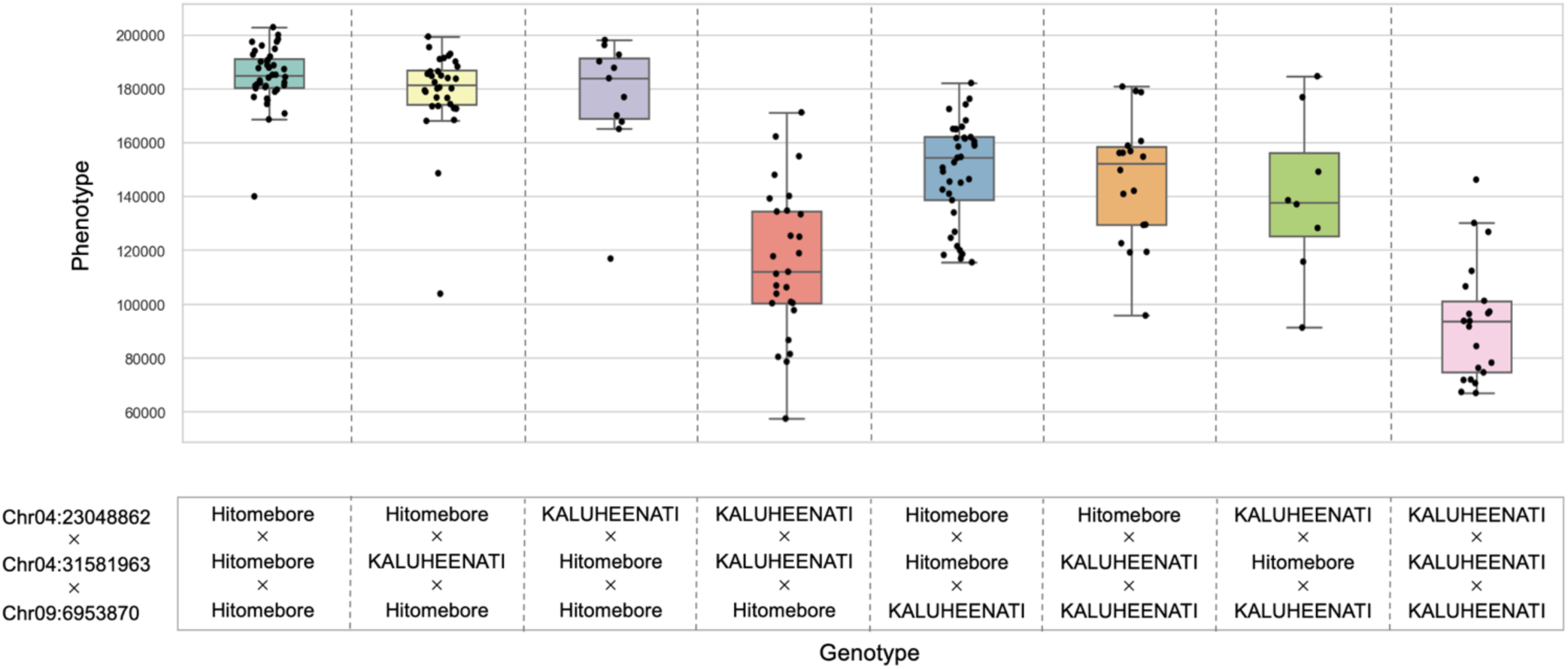
Relationships between rice seed hull color phenotypes and genotypes of the three loci. Boxplots showing the phenotypic values of RILs (Y-axis) separately for the different combinations of genotypes in the three loci (X-axis). When the genotype of chr09:6953870 is KALUHEENATI genotype, phenotypic values are consistently low independent of the genotypes of the other two loci. When the genotypes of chr04:23048862 and chr04:31581963 are both KALUHEENATI types, the phenotypic values tend to be low independent of chr09:6953870.

**Figure S4.**
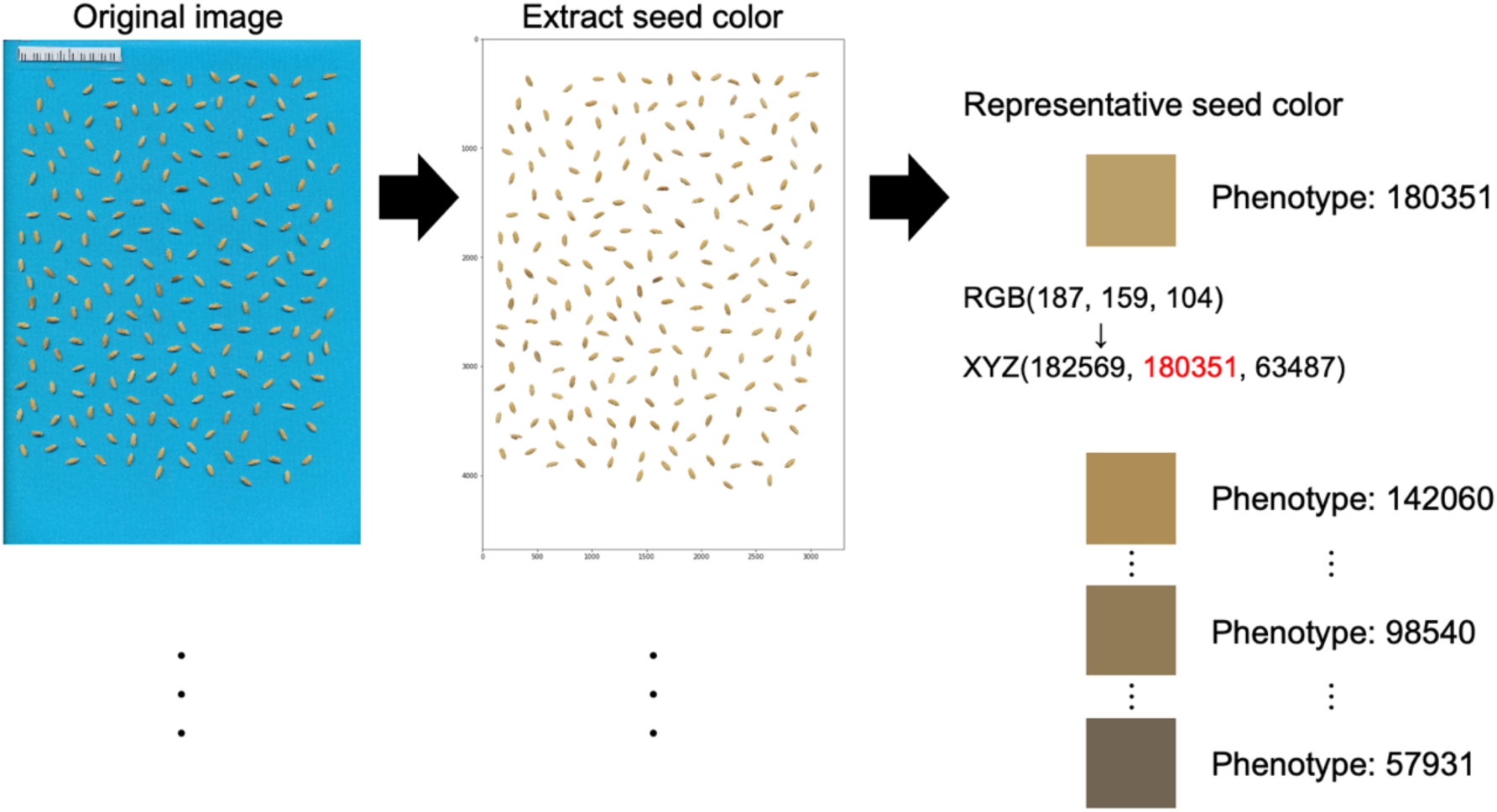
Phenotyping of rice seed hull color. We first trimmed only the image of seeds. Next, we extracted the representative color of the seeds using Principal Component Analysis(PCA). Finally, we converted RGB values of representative seed hull color to Y-axis values of CIE XYZ color space.

