## Supplemental Table 3 for "RIL-StEp: epistasis analysis of recombinant inbred lines (RILs) reveals candidate interacting genes that control rice seed hull color"

**Table S3 Values of RIL-StEp coefficients used for each genotype.**

We included three interaction variables ($\boldsymbol{E}_{\boldsymbol{1}}\boldsymbol{\sim}\boldsymbol{E}_{\boldsymbol{3}}$) to Model2 as variables indicating interactions between the selected two SNPs. We set these values to be 0 when considering interaction between Hitomebore genotype of SNP1 and Hitomebore genotype of SNP2.

| SNP1 | SNP2 | $\boldsymbol{S}_{\boldsymbol{1}}$ | $\boldsymbol{S}_{\boldsymbol{2}}$ | $\boldsymbol{E}_{\boldsymbol{1}}$(H*F) | $\boldsymbol{E}_{\boldsymbol{2}}$(F*H) | $\boldsymbol{E}_{\boldsymbol{3}}$(F*F) |
| --- | --- | --- | --- | --- | --- | --- |
| Hitomebore | Hitomebore | 0 | 0 | 0 | 0 | 0 |
| Hitomebore | Founder | 0 | 1 | 1 | 0 | 0 |
| Founder | Hitomebore | 1 | 0 | 0 | 1 | 0 |
| Founder | Founder | 1 | 1 | 0 | 0 | 1 |
